# Early-life PBDE flame retardant exposures cause neurobehavioral alterations in fish that persist into adulthood and vary by sex and route of exposure

**DOI:** 10.1101/2025.04.28.651140

**Authors:** Nicole McNabb-Kelada, Tara Burke, Saro Jayaraman, Lesley Mills, Ashley De La Torre, Madison Francoeur, Hannah Schraeder, Diane Nacci, Bryan Clark, Andrew Whitehead

**Affiliations:** Department of Environmental Toxicology, University of California Davis, Davis, CA 95616, USA; Atlantic Coastal Environmental Sciences Division, US EPA Office of Research and Development, Narragansett, RI 02882, USA

## Abstract

Developing organisms exhibit varying susceptibility to environmental pollutants depending on the timing of exposure during development. Early-life development is particularly vulnerable, and many marine species spend early life in nearshore environments, elevating their risk of pollutant exposure. Polybrominated diphenyl ether (PBDE) flame retardants are persistent and ubiquitous pollutants, particularly in nearshore marine environments. They disrupt early-life development and pose ongoing risks for ocean health. However, whether early-life exposure influences neurotoxic effects, and whether those effects are durable throughout life, remains poorly understood. Using *Fundulus heteroclitus*, we tested whether early-life exposure to 2,2’,4,4’,5-pentaBDE (BDE-99) leads to persistent behavioral and molecular alterations in adulthood and whether outcomes differ by exposure route. We conducted two complementary experiments comparing progenitor (maternal) exposure and direct waterborne exposure during embryonic development. Similar doses to developing fish were achieved in both experiments.

After hatch, all fish were reared in clean water until adulthood over two years later, at which time we assessed impacts on behavior and brain gene expression. Both exposure routes led to long-term hyperactivity and reduced anxiety-like behavior, but specific effects varied by dose and sex. Progenitor exposure altered behavior and the brain transcriptome in F1 males (females not tested), whereas direct embryonic exposure affected behavior in females and not males.

These findings highlight the importance of maternal influences, such as chemical metabolites, altered lipid provisioning, small molecules, and epigenetic imprinting, alongside chemical transfer, in shaping the long-term persistence of behavioral and molecular effects from early-life exposure. The distinct effects between exposure routes suggest that dosing with chemical alone is not the only determinant of toxicity. We conclude that maternal factors that are modified by exposure significantly contribute to health outcomes in developing offspring, emphasizing the need to consider exposure route when assessing risks from persistent pollutants. Given that maternal and environmental exposures co-occur in nature, future studies should assess their combined impacts to better predict real-world risks.

## Introduction

During different developmental periods, organisms are variably susceptible to environmental factors such as nutritional deficiencies, maternal stress, hypoxia, climate-related changes, and chemical pollutants^1–6^. Exposure during particularly sensitive developmental windows can result in long-term physiological and behavioral consequences, some of which may not become apparent until later in life^7–9^. This phenomenon contributes to the Developmental Origins of Health and Disease (DOHaD) paradigm, which posits that environmental conditions experienced during early development have lasting impacts on health and disease risk^10^. Early life is especially vulnerable because biological systems, including the nervous, endocrine, and immune systems, are still developing, making them highly susceptible to disruption.

Perturbations at this stage can lead to lasting alterations in metabolic programming^11^, immune function^12^, and neurodevelopment^13–15^. Early-life exposure may be particularly relevant for marine species because many start life in nearshore environments^16^ where pollutant exposures are concentrated (compared to the open ocean). Certain environmental toxicants, such as lead^17–19^, methyl mercury^20,21^, bisphenol A (BPA)^22,23^, phthalates^24^, dioxins^25^, polycyclic aromatic hydrocarbons (PAHs)^26,27^, and polychlorinated biphenyls (PCBs)^28,29^, have been linked to adverse outcomes following early-life exposure, with some manifesting later in life. These environmental exposures during critical periods can “reprogram” an organism’s physiology, leading to persistent dysfunctions that are not always immediately detectable. Despite this, many toxicological assessments focus primarily on acute effects. More research is needed to understand the chronic and lifelong consequences of exposure to environmental toxicants to better inform ecotoxicological risk and impact assessment.

One class of environmental pollutants that interferes with early development is polybrominated diphenyl ethers (PBDEs). PBDEs are brominated flame retardants that were extensively used in consumer products and readily leach out of products into the environment^30^. Though most were phased out globally in the early 2000s, PBDEs continue to be released into the environment through product disposal and recycling^31–34^, and some are still produced in Asia^35,36^. Since PBDEs are highly lipophilic and persistent, they bioaccumulate in tissues and biomagnify through food webs^37,38^. PBDEs are therefore ubiquitous in the environment and in terrestrial and aquatic species^37–44^, including marine species commonly consumed by people. A major concern with PBDE exposure is its ability to transfer from parent to offspring during early development. In humans, PBDEs have been detected in placenta, umbilical cord blood, and breast milk, confirming prenatal and postnatal exposure pathways^45–52^. In oviparous species such as fish, PBDEs can also be deposited into eggs, leading to early-life exposure via maternal transfer^53,54^. Despite the substantial reduction in use, PBDEs remain pervasive in the environment, posing ongoing risks to developing organisms. This raises critical concerns about the persistence of PBDE-induced developmental effects and how exposure route (maternal vs. environmental) modulates long-term outcomes.

The route through which developing embryos are exposed to pollutants like PBDEs, whether via maternal transfer or direct environmental exposure, may significantly influence toxicity. Maternal transfer introduces not only the toxicant itself but also its metabolites^55–57^, along with maternal hormones, immune factors, and other bioactive molecules that shape early development and may have been perturbed by maternal exposure. Maternal exposure may also alter nutrient allocation^58^, affecting key resources such as lipids, antioxidants, and immune-related molecules^59,60^. These changes, along with disruptions in hormone levels and hormone transfer^61–63^, can modify developmental trajectories and influence offspring resilience. In addition to these maternal effects, parental exposure can induce stable epigenetic modifications that cause long-term changes in gene expression^64,65^. In contrast, direct embryonic exposure involves uptake of the toxicant from the surrounding environment without the maternal physiological or parental epigenetic influences, meaning toxicity is driven primarily by the chemical itself. These differences underscore the need to investigate whether and how exposure route influences the persistence and nature of PBDE-induced developmental effects. To address these uncertainties, we designed experiments to test whether early-life exposure to PBDEs leads to persistent effects in adulthood and whether the exposure route (maternal transfer vs. direct embryonic) alters these outcomes.

The developing brain is especially sensitive to perturbation, meaning that even low levels of toxicant exposure during early development can lead to persistent impairments in cognitive functions and various behavioral alterations^1,13^ that influence fitness. Indeed, exposure to PBDEs has been linked to developmental neurotoxicity, where early-life exposure disrupts brain development through mechanisms including thyroid hormone dysregulation, oxidative stress, and interference with calcium signaling^66,67^. Epidemiological studies have associated prenatal and childhood PBDE exposure with impaired motor skills, cognitive deficits, and lower IQ^14,68^. Among the PBDE congeners, 2,2’,4,4’,5-pentabromodiphenyl ether (BDE-99) is one of the most prevalent in human and wildlife samples, including marine fish^69–71^. In mammals, both pre-and postnatal exposures to BDE-99 affect spontaneous locomotor activity, habituation to anxiety-promoting situations, and learning abilities^67,72,73^. However, not all studies have examined whether these changes persist into adulthood, and those that have found mixed results. The role of exposure route in shaping these long-term effects remains unclear, yet it may be a critical factor influencing neurodevelopmental outcomes. This study, therefore, aimed to test whether early-life exposure to BDE-99 results in persistent behavioral effects and molecular alterations in the brain, where we examined outcomes following exposure via maternal transfer compared to outcomes following direct aqueous exposure.

For our experiments, we used Atlantic killifish (*Fundulus heteroclitus*), a well-established model in ecotoxicology^74^. This nonmigratory marine fish inhabits salt marsh estuaries along the U.S. Atlantic coast^75^, including highly polluted environments^76^. Killifish harbor high levels of genetic diversity^77,78^, allowing us to capture interindividual variability in responses to toxicant exposures that may be more representative of the variance expected from wild marine species, compared to more typical laboratory-bred model organisms.

Neurodevelopmental pathways are highly evolutionarily conserved across vertebrates^79^, such that findings in *F. heteroclitus* should be of broad relevance for other marine and terrestrial vertebrates.

This study investigated the long-term effects of early-life exposure to BDE-99 in *F. heteroclitus*, testing whether brief exposures during embryo development (prior to hatch) lead to persistent behavioral changes and alterations in brain gene expression in adulthood. In a companion study we focused on effects that propagate across generations (McNabb-Kelada et al., unpublished), whereas in this study we focused on long-term within-generation effects. In this study we specifically compared two exposure routes during embryonic development: progenitor (maternal) exposure and direct waterborne exposure. In the progenitor exposure route, adult fish were fed control or BDE-99-contaminated diets for 64 days during the spawning season, leading to maternal deposition of ingested chemical into their eggs and exposure of F1 offspring starting from the earliest stages of development. In the direct exposure route, embryos from unexposed parents were directly exposed to BDE-99 in water from 1-7 days post-fertilization (dpf). After hatch, all embryos were raised in clean water until reaching adulthood. Over two years after exposure, we assessed anxiety-like behavior and conducted whole-brain transcriptomic analyses to determine whether early-life exposure to BDE-99 resulted in persistent neurobiological effects. This approach was designed to test our hypotheses that (1) exposure to BDE-99 during short, discreet windows during early life affects brain developmental trajectories such that perturbations are persistent throughout life; and (2) the two exposure routes (maternal vs. direct embryonic) would yield different outcomes, revealing distinct mechanisms of toxicity associated with maternal transfer versus direct exposure. By addressing these hypotheses, this study provides insight into how early-life exposure to PBDEs affects long-term neurobehavioral and molecular outcomes and highlights the potential for distinct mechanisms contributing to toxicity between maternal and environmental exposure routes.

## Methods

We conducted two complementary experiments to test for the long-term effects of early-life exposure to BDE-99 on adult behavior, where we contrasted effects that emerged from two different routes of exposure (Figure 1). In Experiment 1, we investigated the long-term effects of early-life exposure to BDE-99 via maternal transfer by feeding adult progenitors contaminated diets and assessing behavioral, reproductive, and molecular outcomes in their offspring (F1 generation) during adulthood. In Experiment 2, we examined the long-term effects of direct embryonic (waterborne) exposure to BDE-99 by evaluating the same behavioral, reproductive, and molecular responses in adulthood. Though many of the methods were identical between experiments, we describe them sequentially below.

**Figure 1:**
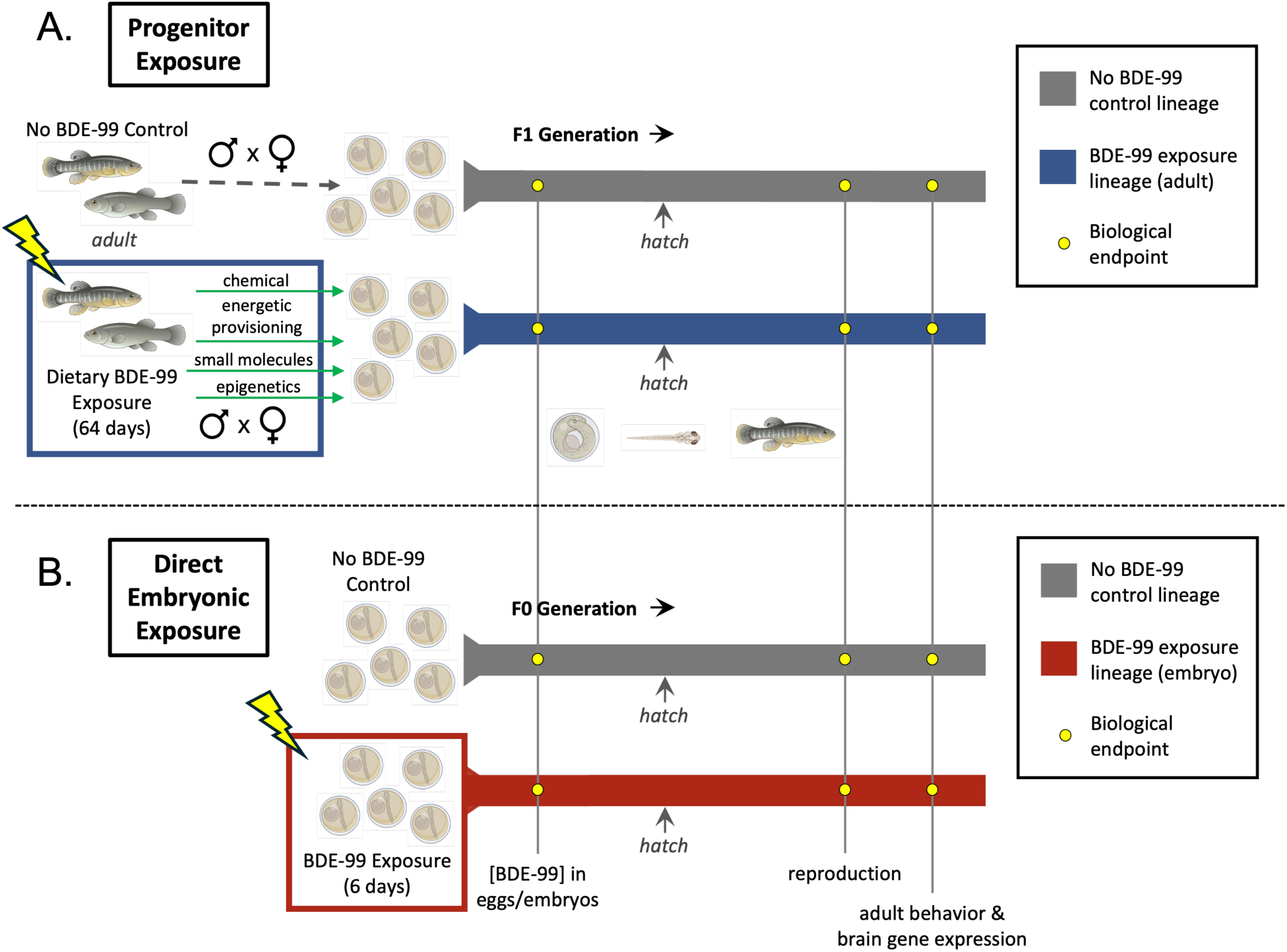
Experimental design overview for two complementary experiments to test for persistent effects of early-life exposures to BDE-99 via progenitor (A, Experiment 1) and direct embryonic (B, Experiment 2) routes. Developing killifish embryos were exposed to BDE-99 either through (A) maternal transfer from adult progenitors or (B) direct waterborne exposure. A) Progenitor Exposure: Adult F0 fish were exposed to control or BDE-99-contaminated (blue box) diets for 64 days during the breeding season. First-generation (F1) descendants (gray line = control, blue line = progenitor BDE-99 exposure) were produced by mating males and females within each lineage and were raised in uncontaminated conditions until adulthood. B) Direct Embryonic Exposure: Embryos from unexposed parents were exposed to either control or BDE-99 (red box) via waterborne exposure from 1-7 days post-fertilization (dpf). These (F0) fish were then raised in uncontaminated conditions until adulthood (gray line = control, red line = direct embryonic BDE-99 exposure). Biological endpoints, including fertilization success, anxiety-like behavior, and brain gene expression, were measured at the same time points in adulthood across both exposure experiments. Fish and embryo illustrations sourced from BioRender.com.

### Experiment 1: Progenitor Exposure Chronological Overview

During the breeding season, adult wild-caught killifish were fed either control or BDE-99-contaminated diets for 64 days. This exposure resulted in parental (maternal) transfer of BDE-99 to their offspring (F1 generation). F1 fish were then reared in clean water until adulthood (2 years post-hatch) when reproductive endpoints were measured (2 years post-hatch) (Figure 1A) and behavior and brain gene expression measured (2.5 years post-hatch).

### Adult Fish Collection and Husbandry

Adult Atlantic killifish (∼10 g wet weight (ww)) were collected from Jerusalem, Narragansett, RI, USA (41°22’58.4” N, 71°31’09.6” W)^80^ then housed at the US Environmental Protection Agency (EPA) laboratory in Narragansett in large (300 L, 80 gal) tanks (continuous flow, 23°C, 5 µm-filtered, clean seawater, 14:10 h light:dark cycle, fed TetraMin Tropical Flakes *ad libitum*). After one month of acclimation, fish were individually PIT tagged and distributed into 38-L (10-gal) tanks. Each treatment included three replicate tanks, each containing 14 fish (8 females, 6 males). All procedures were conducted following EPA Animal Care and Use Protocols (IACUC, ACUP # Eco23-03-002, Eco23-11-001, and Eco23-07-001).

### Diet Preparation for Progenitor BDE-99 Exposure

BDE-99 (2,2’,4,4’,5-pentabromodiphenyl ether; >99% pure) was from AccuStandard (Lot no. 27669; New Haven, CT, USA). For the BDE-99 stock solution (2.5 mg/mL), 10 mg of neat BDE-99 was dissolved in 4 mL of acetone. For diet preparation, two dosing solutions were made (1.4 mL each): BDE-99_MED and BDE-99_HI. BDE-99_MED contained 52 µL of the stock solution and 1348 µL of acetone. BDE-99_HI contained 209 µL of the stock solution and 1191 µL of acetone. Each batch of diet consisted of 40 g of ground TetraMin Tropical Flakes, amended with either 1.4 mL of acetone (carrier control) or BDE-99 dosing solution. To supplement the diet, a nutritional mixture was prepared by blending chopped spinach, brine shrimp, krill, mackerel, and sardines. Each batch of diet was supplemented with 450 g of the nutritional mixture, which was consolidated using gelatin.

### Progenitor Dietary Exposure to BDE-99

Fish were fed once daily either an acetone-amended control or a BDE-99-contaminated diet (∼15% of body weight per day) for 64 days (July 9 – September 11, 2019; treatment diets were provided five days per week, and control diet the remaining two days per week). Fish were exposed to one of three dietary treatments: control, BDE-99_MED (37.5 ng/g fish ww/day), or BDE-99_HI (150 ng/g fish ww/day). Target internal doses of BDE-99 were 1 and 4 μg/g fish dw.

### Spawning

Fish were manually strip-spawned five times throughout the exposure period to produce F1 offspring that were developmentally exposed to BDE-99 (or control) via maternal transfer. Eggs and milt collected from fish within the same tank were pooled for fertilization but kept separate between tanks. Embryos were incubated at 23°C under a 14:10 h light:dark cycle. At 4 dpf, unfertilized eggs were flash-frozen and stored at-80°C for BDE-99 quantification. For two years, F1 fish were raised in clean water, then spawned to assess impacts of progenitor exposure on reproduction. Due to low female numbers during the 2021 spawning season in both F1 control and BDE-99 treatments, males were mated with external control females. Eggs from 20-36 females were pooled, divided into approximately equal groups, and fertilized with pooled milt from males within each treatment. The fertilization rate was recorded at 4 dpf for each spawning time point. After confirming normality of data (Shapiro-Wilk test), one-way Analysis of Variance (ANOVA) was used to test for treatment effects on fertilization rate (percentage of fertilized eggs per treatment). Statistical analyses were conducted using GraphPad Prism version 10.4.1 for macOS.

### Chemical Analysis

Accelerated Solvent Extraction (ASE 200; Dionex Corporation, Sunnyvale, CA, USA) was used to extract BDE-99 from progenitor treatment diets, adult fish following dietary exposure, F1 eggs from progenitor exposures, and F0 embryos from direct embryonic exposure^81^. Eggs (40-150 mg ww) were ground with a mortar and pestle, spiked with IS, combined with acetone, vortexed for 30 seconds, and centrifuged at 2000 rpm for 5 minutes. Extracts were back-extracted with hexane and concentrated using ultra-high-purity nitrogen. BDE-99 concentrations were measured using Gas Chromatography-Mass Spectrometry (GC-MS) on an Agilent Technologies (Santa Clara, CA, USA) 6890N GC equipped with a 5973 mass selective detector and an Agilent J&W DB-5ms Ultra Inert analytical column (15 m x 0.25 mm x 0.25 µm). Helium was used as the carrier gas at a constant flow rate of 1.6 mL/min. Calibration standards for BDE-99 (50-2500 ng/µL) were prepared in heptane, and a seven-point calibration curve was generated for quantification. Data processing was performed using ChemStation software, and BDE-99 concentrations were reported on a wet weight (ww), dry weight (dw), and lipid weight basis. To ensure quality assurance and quality control (QA/QC), procedural blanks, standard reference material (SRM 1947, Lake Michigan fish tissue), analytical duplicates, and matrix spikes were routinely processed. Procedural blanks showed no contamination, and the laboratory-measured BDE-99 concentration in SRM 1947 was within 10% of the NIST-certified reference value. Analytical duplicates exhibited a relative standard deviation (RSD) of <7%, while matrix spike recoveries ranged from 90% to 104%, confirming method accuracy and precision.

### Rearing of Progenitor Exposure F1 Fish

F1 embryos from F0 progenitor exposed and control lineages across multiple spawning time points were incubated in clean seawater (23°C, 14:10 h light:dark cycle). At 7 dpf, embryos were transferred to individual wells in 12-well tissue culture plates. At 10 dpf (late organogenesis, stage 34 ^82^), embryos were screened for the presence of developmental abnormalities, including pericardial edema, heart abnormalities, cranial or caudal hemorrhaging, and reduced body or head size^83^. There were no treatment effects on developmental abnormalities. At 14 dpf, plates were rocked to stimulate hatching, after which larvae were fed *Artemia ad libitum*. Larvae were transferred to 9.5 L (2.5 gal) tanks after 14 dph, then gradually transitioned to 19 L (5 gal) and later 38 L (10 gal) tanks to accommodate growth. Juvenile fish were housed at 3-10 fish per gallon.

### Novel Tank Diving Test

At 2.5 years post-hatch, a novel tank diving test was conducted to assess anxiety-like behavior in adult male fish, as low female numbers across all treatments precluded their inclusion. The experimental setup followed the protocol reported by Levin et al. 2007, with modifications^84^. Two adjacent rectangular tanks (33.6 cm W x 33.9 cm H x 8.2 cm D) were each filled with ∼4.5 L seawater (21.5-23°C). An LED light pad (Huion, Shenzhen, China) provided uniform overhead illumination. Light levels (180 ± 10 lx). A Basler acA1300-60gmNIR camera was positioned 120 cm from the front of the tanks to record the behavioral trials. Video data were captured (30 frames per second) and analyzed using EthoVision XT software (version 11.5.1026, Noldus, Wageningen, Netherlands) for fish tracking and behavioral assessment.

At the start of each trial, fish were released into the novel tank environment and recorded for 10 minutes. Anxiety-like behavior was evaluated using *total distance moved* (cm per 10 min), *duration spent in the top zone* (top half of the tank; percent time), *number of transitions to the top zone*, and *latency to enter the top zone* (first entry; seconds). After trials, brains were preserved for transcriptomics. Behavioral data were not normally distributed (Shapiro-Wilk test), such that the Kruskal-Wallis test was used to test for treatment effects, followed by Dunn’s post-hoc tests. Statistical analyses were conducted with GraphPad Prism version 10.4.1.

### RNA-Seq and Read Count Quantification

Brains were stored at-80°C until homogenization using 2.8 mm ceramic beads. Lysates were suspended in Lysis Binding Buffer^85^. After centrifugation, messenger RNA (mRNA) was extracted from the supernatant using oligo (dT)25 Dynabeads^TM^ mRNA DIRECT^TM^ Purification Kit. The Breath Adapter Directional sequencing (BrAD-seq) method^85^ was used to prepare strand-specific RNA-seq libraries (13 PCR cycles), with fragment priming using a random hexamer (GTGACTGGAGTTCAGACGTGTGCTCTTCCGATCTNNNNNNNN). Following Qubit^TM^ quantification (dsDNA High Sensitivity Quantification Assay Kit), uniquely indexed individual libraries were pooled and then sequenced across two lanes of NovaSeq X Plus PE150 (Illumina, San Diego, CA, USA) by IDseq Inc. (Davis, CA, USA). Each sample yielded ∼13 million raw reads. Eight biological replicates were included for each of three treatment groups (control and the two highest BDE-99 progenitor exposure lineages (2.00 and 3.50 µg/g); 24 samples total). Raw sequencing reads were processed using a Snakemake pipeline (https://github.com/JoannaGriffiths/RNASeq-snakemake-pipeline), which included quality-checking (FastQC^86^), trimming (fastp^87^) to remove adapter sequences and low-quality bases, and Salmon^88^ to map processed reads to the *F. heteroclitus* reference genome (GenBank MU-UCD_Fhet_4.1)^89^ and for transcript-level quantification. Raw read counts were then summed to the gene level using gene annotation data for gene-level analyses.

### WGCNA and Functional Enrichment

We used weighted gene correlation network analysis (WGCNA)^90^ to test for progenitor exposure effects on brain gene expression. A variance-stabilizing transformation (VST) in DESeq2 ^91^ normalized for variation in library size. The top 10% most variable genes (n=3,164) were selected, as recommended for WGCNA, then pairwise gene expression correlations were computed to construct a signed adjacency matrix. Optimal soft-thresholding power was selected using the scale-free topology criterion. Co-expressed gene expression modules were identified using hierarchical clustering with dynamic tree cutting. Module eigengene values (the first principal component of module expression) were extracted to summarize expression patterns across samples. Eigengene values were tested for significant associations with treatment (developmental BDE-99 exposure) using one-way ANOVA. The dataset included three treatments (CTL, 2.00 µg/g BDE-99 dose, 3.50 µg/g BDE-99 dose), including 8 biological replicates per treatment, totaling 24 samples. Modules showing significant treatment effects (FDR-adjusted p<0.05) were tested for Gene Ontology (GO) enrichment (Mann-Whitney *U* test^92^). All analyses were conducted in R (version 4.4.2)^93^ and RStudio (version 2024.09.1+394).

### Experiment 2: Direct Embryonic Exposure Chronological Overview

Fish embryos from unexposed parents were exposed to either control conditions or BDE-99 from their surrounding water from 1-7 dpf (Figure 1B). These directly exposed F0 embryos were then reared to adulthood in clean water. Experimental endpoints measured in adults were the same as those measured in the progenitor exposure (Experiment 1): adult reproductive assessment of fertilization success at 2 years post-hatch, adult behavior (novel tank diving test) at 2.5 years post-hatch, and brain transcriptomics.

### Embryo Collection and Fertilization

Embryos were obtained through mass manual spawning of lab-bred adult fish from a population originally collected from Scorton Creek, Sandwich, MA (41°45’53.6” N, 70°28’48.0” W)^94–97^, that had always been maintained under clean conditions. Spawning was conducted with four separate breeding tanks, with embryos from each tank kept separate. In each tank, eggs were collected from 20-30 females, pooled, then fertilized with pooled milt from 20-30 males from the same tank.

### Direct Embryonic Exposure to BDE-99

A BDE-99 stock solution (2.5 mg/mL) was made by mixing 10 mg of neat BDE-99 into 4 mL of acetone. Fertilized embryos from the four breeding groups were equally distributed among three treatment conditions (control and two BDE-99 concentrations) and further subdivided into two replicates per treatment, resulting in 24 exposure jars (4 breeding groups x 3 treatments x 2 replicates per treatment). Embryos were placed in glass jars (1 embryo per 2 mL) and exposed from 1-7 dpf. Exposure solutions contained a 1% dosing solution, consisting of either carrier (acetone, control) or BDE-99 in acetone at final concentrations of 6.2 ng/mL (MED) and 26 ng/mL (HI). At 7 dpf, embryos were transferred to clean water and reared as described for Experiment 1. Embryos from one of the breeding groups were archived for BDE-99 quantification.

### Experimental Endpoints

To assess the long-term reproductive effects of early-life exposure to BDE-99, males and females were crossed with mates from within their treatment lineages. Eggs were collected via manual strip-spawning, pooled, and fertilized with pooled milt from males in a different replicate group within the same exposure condition (to minimize mating with close relatives). One-way ANOVA was used to test for treatment effects on fertilization rate (percentage of fertilized eggs per treatment at 4 dpf) following confirmation of normality (Shapiro-Wilk test).

At 2.5 years post-hatch, the novel tank diving test was conducted to assess anxiety-like behavior in both male and female fish, as described for Experiment 1. A generalized linear model (GLM) with least squares regression was used to test the effects of sex and BDE-99 dose on behavioral outcomes. The model included the main effects of sex (categorical: male, female) and dose (categorical: CTL, MED, and HI (0, 1.00, and 2.70 µg/g)), and their interaction. GLMs were chosen because data did not meet normality and homoscedasticity assumptions (Shapiro-Wilk test, Spearman’s test). To correct for heteroscedasticity, inverse weighting (1/Y) was applied. All statistical analyses were performed in GraphPad Prism version 10.4.1.

### RNA-Seq and WGCNA

Brains from adult fish that had been exposed during embryogenesis were processed for transcriptomics at the same time and using the same methods as described for Experiment 1 samples. Each sample yielded ∼12 million raw reads. The dataset included two treatments (CTL and HI BDE-99 dose), with both combined and separate analyses for males and females. For males, there were 2-6 replicates each from 3 exposure replicates per treatment, totaling 27 samples. For females, there were 3-6 replicates each from 3 exposure replicates per treatment, totaling 30 samples. Raw sequencing reads were quality-checked and processed as previously described. WGCNA was performed using the same parameters and workflow as described for Experiment 1.

## Results

### Dosing

Developmental exposures to BDE-99 achieved our targeted doses of 1 to 4 µg/g. The progenitor exposure (Experiment 1) achieved a five-dose range spanning from 0.47 to 3.50 µg/g dw (Figure 2A), and the direct embryonic exposure (Experiment 2) achieved an overlapping two-dose range from 1.00 to 2.70 µg/g.

**Figure 2:**
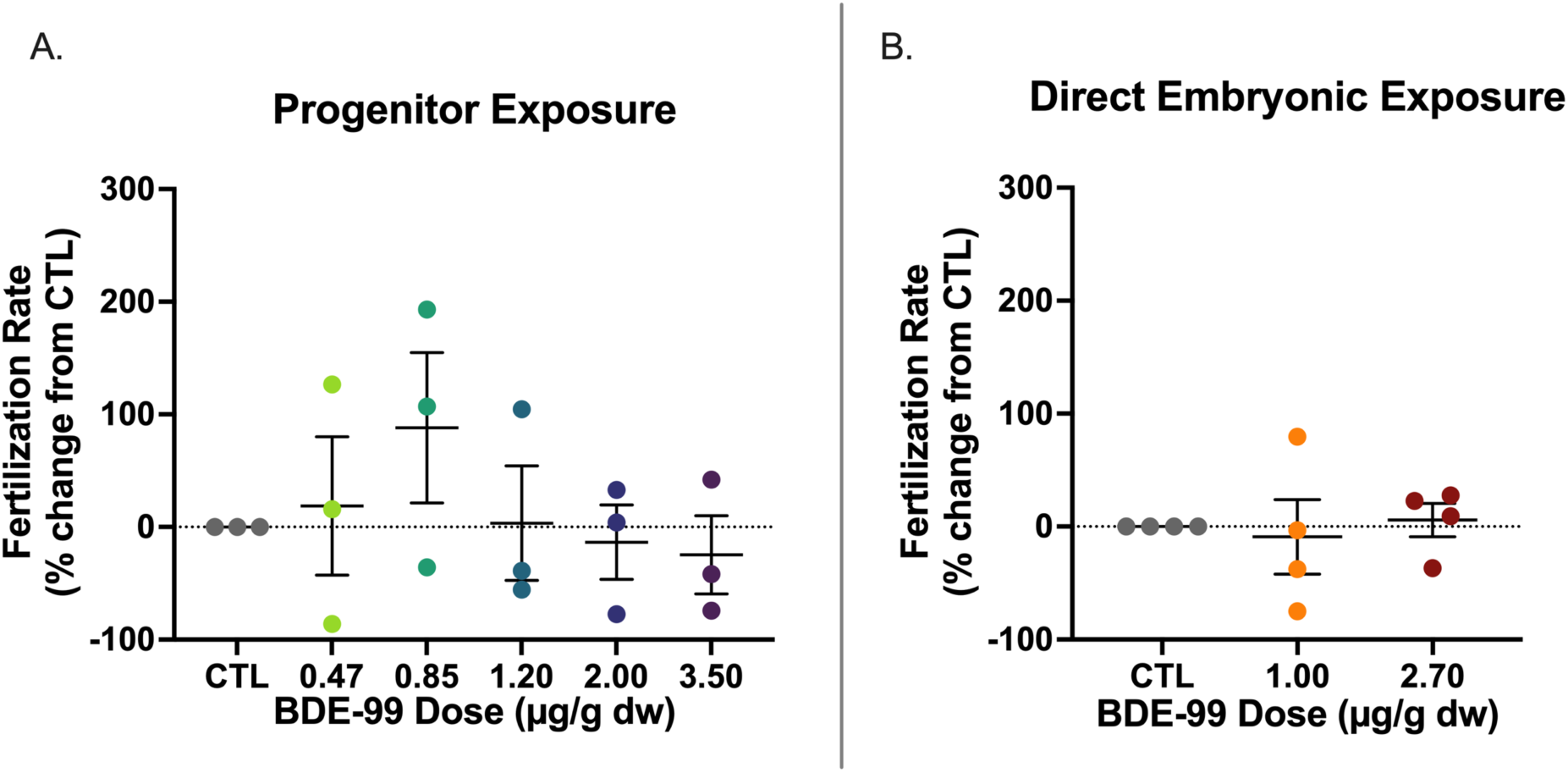
Fertilization rate for adult fish that had been exposed to BDE-99 during early development via progenitor exposure (A) or direct waterborne embryonic exposure (B). Fertilization rate was calculated as the percentage of eggs fertilized successfully at each spawning time point and is expressed as the percent change from the control (CTL) mean. A) At 2 years post-hatch, males from the progenitor exposure experiment were crossed with external control females over three spawning time points. Male fish originated from exposure groups with the following BDE-99 doses measured in eggs: 0 (CTL), 0.47, 0.85, 1.20, 2.00, and 3.50 µg/g dry weight (dw). B) In the direct embryonic exposure experiment, males and females (2 years post-hatch) were crossed within their respective control or BDE-99 treatments over four spawning time points. Fish originated from exposure groups with the following BDE-99 doses measured in subsets of embryos: 0 (CTL), 1.00, and 2.70 µg/g dw. No statistically significant treatment effects were detected (ANOVA). Error bars represent standard error of the mean (SEM).

### Fertilization Rate Across Both Exposure Routes

Developmental exposure to BDE-99 did not affect later-life fertilization success as adults.

This was consistent whether fish had been exposed through their progenitor (Experiment 1, Figure 2A) or exposed directly (Experiment 2, Figure 2B).

### Progenitor Exposure (Experiment 1): Anxiety-Like Behavior Effects in Adult Offspring

Progenitor exposure to BDE-99 significantly altered multiple dimensions of anxiety-like behavior in adult male offspring, including *total distance moved* (p=0.0002) and *latency to enter the top zone* of the novel tank (p=0.0052). Fish from the 0.85 and 2.00 µg/g groups were hyperactive compared to controls (p=0.0017 and 0.0027, respectively) (Figure 3A). Similarly, fish from these same exposure groups entered the top zone faster than controls (p=0.0168 and 0.0279, respectively) (Figure 3D). The amount of time spent in the top zone was also affected by BDE-99 dose (p=0.0137), with fish from the 2.00 µg/g group spending significantly more time in the top zone relative to controls (p=0.0297) (Figure 3B). Additionally, BDE-99 exposure influenced the *number of transitions to the top zone* (p=0.0035), with fish in the 0.47, 0.85, and 2.00 µg/g exposure groups making significantly more transitions compared to controls (p=0.0296, 0.0097, and 0.0195, respectively) (Figure 3C).

**Figure 3:**
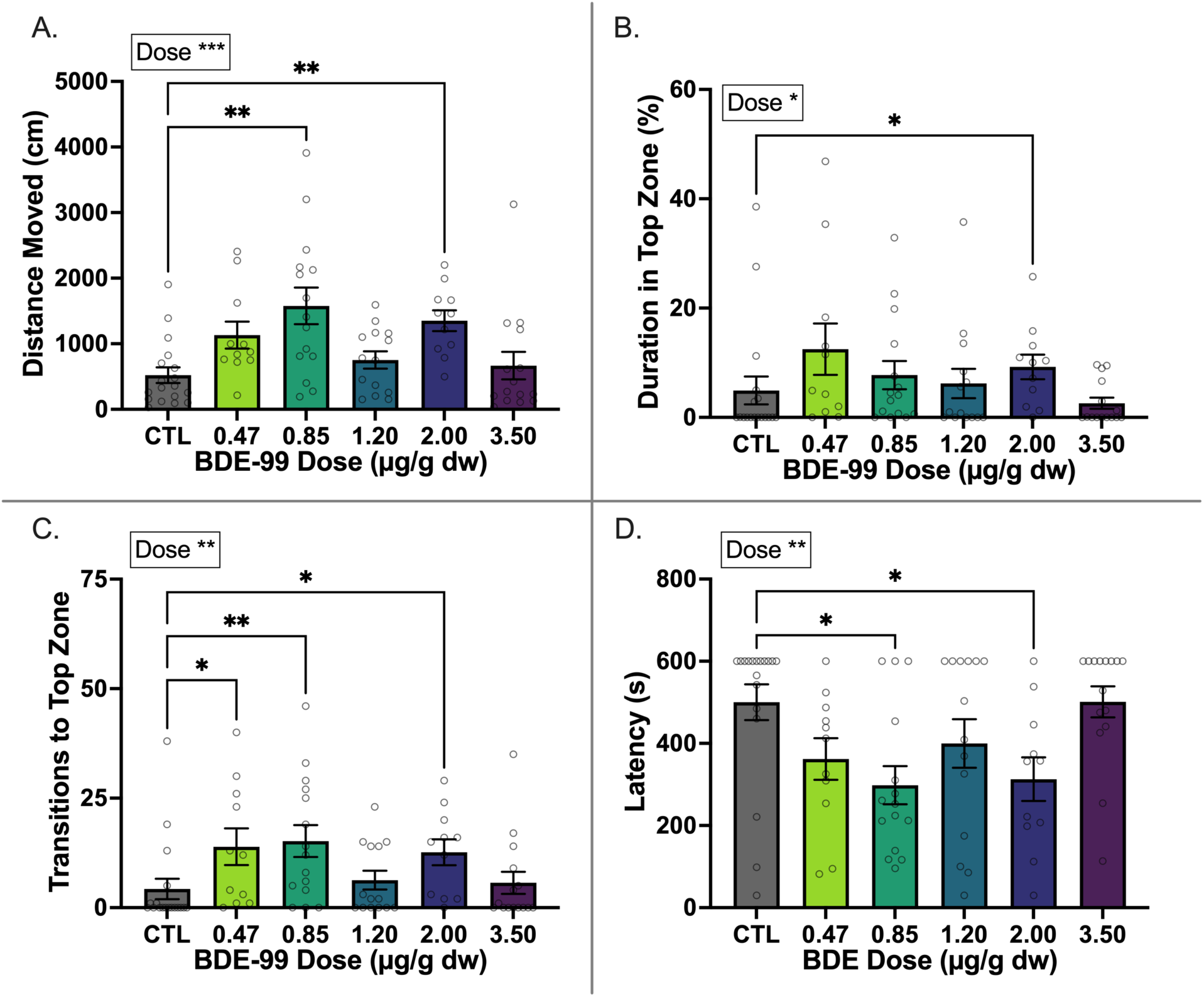
Anxiety-like behavior in adult male fish that were developmentally exposed to BDE-99 via maternal transfer (progenitor exposure). Activity was tracked during 10-min novel tank diving tests at 2.5 years post-hatch. For analysis, the tank was divided into two zones (*top, bottom*). Variables measured include total distance moved (cm) (A), duration (% of time) spent in the top zone of the tank (B), number of transitions to the top zone (C), and latency (seconds) to enter the top zone (D). Fish that never entered the top zone were assigned a maximum latency of 600 seconds (assay duration). Fish originated from exposure groups with the following BDE-99 doses measured in eggs: 0 (control, CTL; n=18), 0.47 (n=11), 0.85 (n=15), 1.20 (n=14), 2.00 (n=11), and 3.50 (n=15) µg/g dry weight (dw). Within a plot, the statistical test outcome (p-value) is shown for the main effect of BDE-99 dose. p<0.05, p<0.01, and p<0.001 are indicated by *, **, and ***, respectively. Brackets with asterisks indicate significant differences from CTL following post-hoc tests. Error bars represent standard error of the mean (SEM).

The 2.00 µg/g exposure group exhibited abnormal activity for all four behavior variables, suggesting that this dose induced the strongest persistent behavioral response. The 0.85 µg/g group showed significant differences in 3 out of 4 variables, while the 0.47 µg/g group exhibited an effect in 1 out of 4 variables. In contrast, fish developmentally exposed to 1.20 or 3.50 µg/g BDE-99 did not differ significantly from controls in any measured behavior (0 out of 4 variables). Overall, we conclude that progenitor exposure to BDE-99 altered anxiety-like behavior in adult descendants in a manner that varied among doses, displaying a non-monotonic dose response, where low to intermediate doses elicited more pronounced and persistent effects than the highest dose.

### Progenitor Exposure: Brain Transcriptome Responses in Adult Offspring

Progenitor exposure to BDE-99 resulted in alterations in F1 offspring brain gene expression that were detectable in adulthood after 2+ years of living in clean conditions. WGCNA identified 14 co-expressed gene modules, each containing between 46 and 611 genes. Of these, three modules (Green, Black, and Magenta) were significantly associated with BDE-99 exposure (adjusted p<0.05), with the Green module (186 genes) containing functionally enriched genes. Principal Component Analysis (PCA) of gene expression in the Green module revealed clear separation between BDE-99-exposed fish and controls, with samples from both BDE-99 exposure groups (2.00 and 3.50 µg/g dw in eggs) clustering together and exhibiting transcriptional divergence from control fish (Figure 4A). Functional enrichment analysis indicated that genes in the Green module were predominantly associated with immune-related pathways (e.g., immune system process, defense response, and response to virus), protein modification pathways (e.g., post-translational protein modification and protein modification by small protein removal), and environmental response pathways (e.g., defense response to other organisms, response to external stimuli, and biological process involved in interspecies interactions) (Figure 4B).

**Figure 4:**
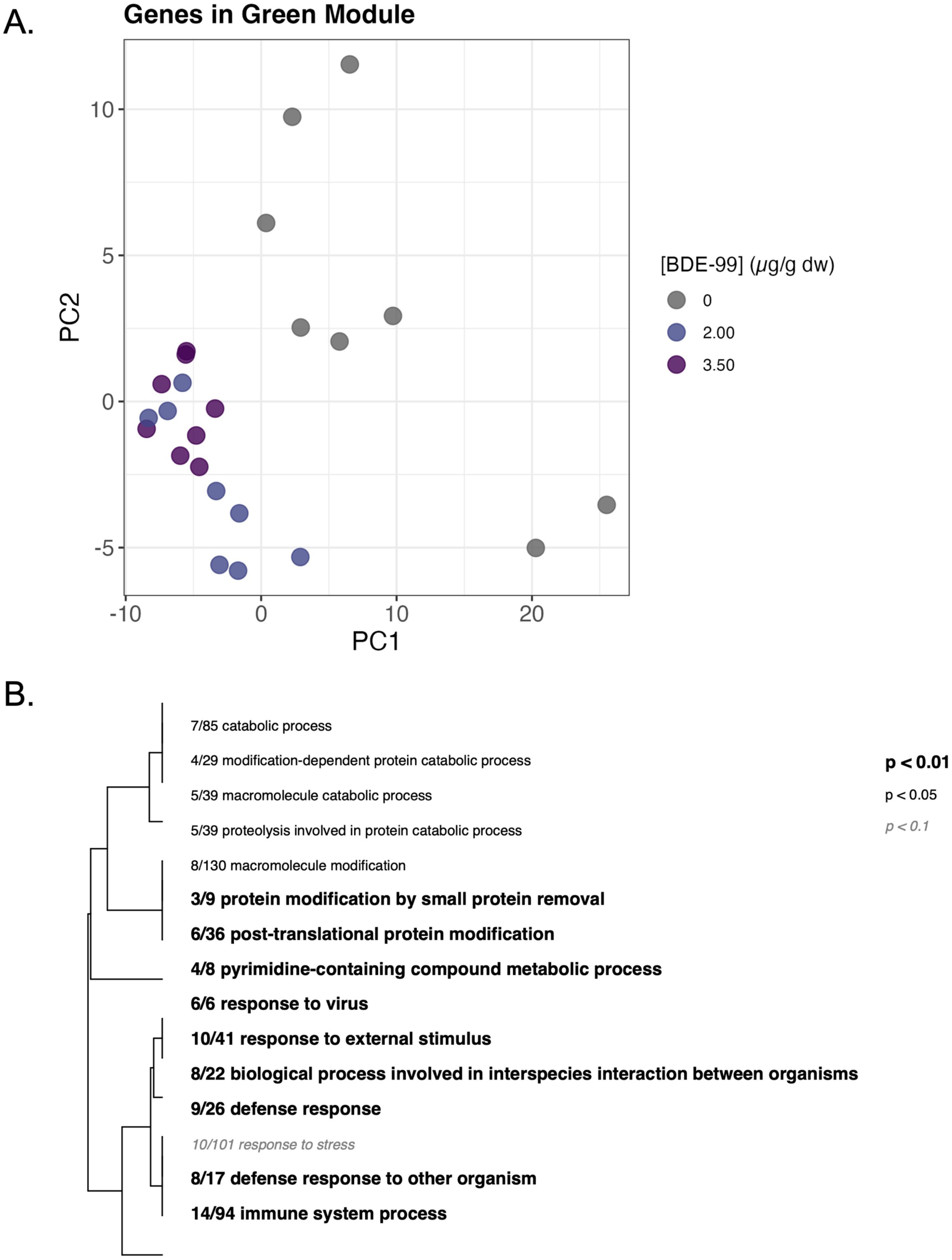
Principal Component Analysis (PCA) of whole-brain gene expression in the Green module and associated Gene Ontology (GO) enrichment analysis in adult fish (2.5 years post-hatch) that were developmentally exposed to BDE-99 via maternal transfer (progenitor exposure). A) PCA plot illustrating variance in gene expression among 186 genes assigned to the Green WGCNA module. Points represent individual samples, colored by BDE-99 exposure group: 0 (control; n=8), 2.00 (n=8), and 3.50 µg/g (n=8). B) Functional enrichment analysis shows GO biological process categories significantly enriched within the Green module. The dendrogram represents hierarchical clustering of enriched GO terms. Fractions indicate the number of genes in the Green module relative to the total number of genes assigned to that category. GO terms are visually distinguished by statistical significance: bolded text for p<0.01, regular text for p<0.05, and italicized text for p<0.1.

### Direct Embryonic Exposure (Experiment 2): Anxiety-Like Behavior Effects in Adults

Early-life exposure to BDE-99 altered multiple dimensions of behavior in adulthood, with effects shaped by interactions between sex and dose. For *total distance moved*, there was a significant dose-by-sex interaction (p=0.0051, Figure 5A). For *number of transitions to the top zone,* there was a significant main effect of dose (p=0.0133) and a dose-by-sex interaction (p=0.0011, Figure 5C). No significant effects of sex, dose, or their interaction were detected for *duration spent in the top zone* (Figure 5B). For *latency to enter the top zone*, a significant dose-by-sex interaction was observed (p=0.0342), but no main effects of sex or dose were observed (Figure 5D). These results indicate that direct embryonic exposure to BDE-99 during development manifests in later life (adults 2+ years later) as perturbations of locomotor activity and anxiety-like behaviors in a sex-dependent manner. Dose-dependent effects were primarily observed in females.

**Figure 5:**
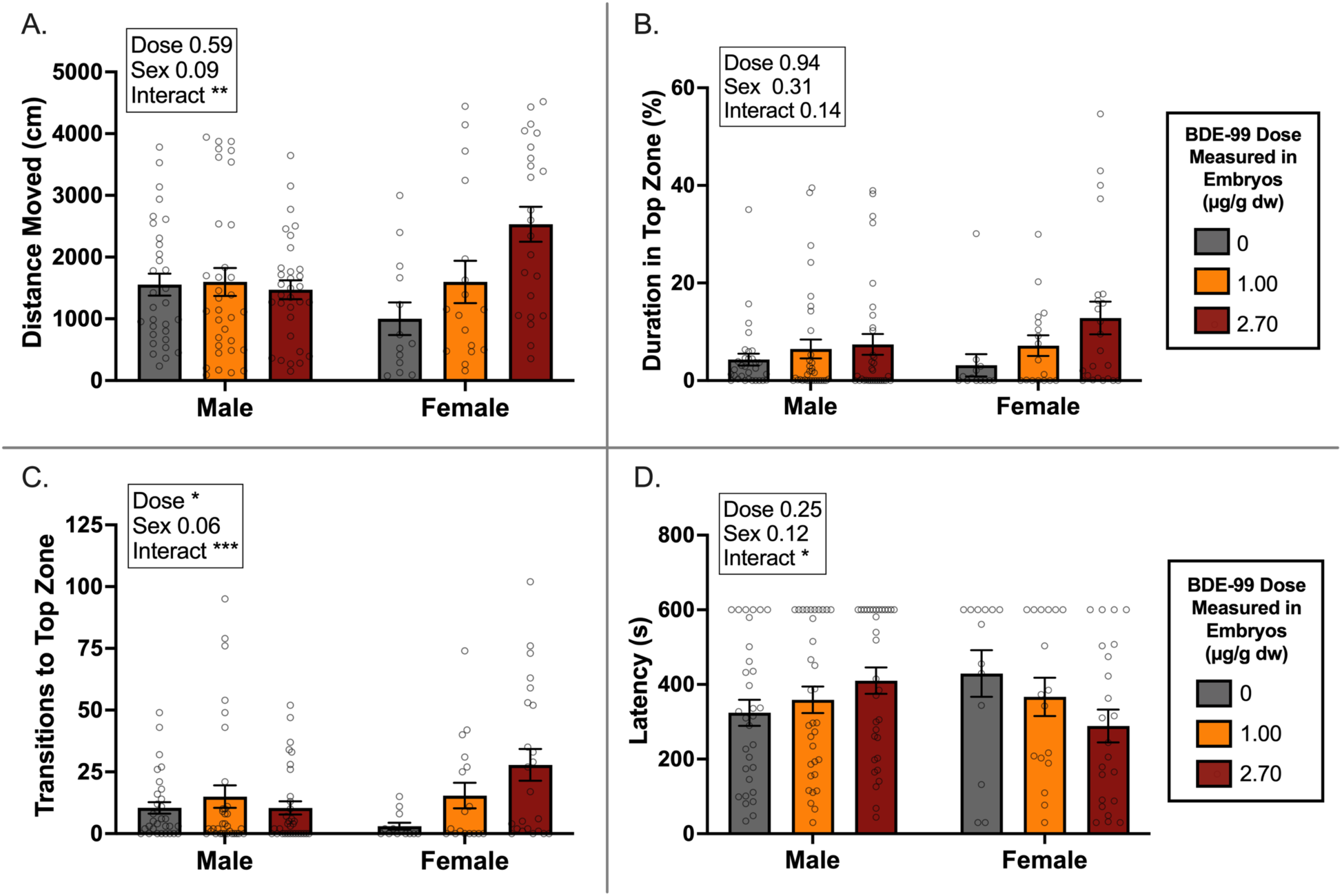
Anxiety-like behavior in adult fish following direct embryonic exposure to BDE-99. Fish were exposed to control or BDE-99 waterborne treatments from 1-7 days post-fertilization (dpf), and activity was tracked during 10-min novel tank diving tests at 2.5 years post-hatch. Data for males and females are shown separately within each plot. For analysis, the tank was divided into two zones (*top, bottom*). Variables measured include total distance moved (cm) (A), duration (% of time) spent in the top zone of the tank (B), number of transitions to the top zone (C), and latency (seconds) to enter the top zone (D). Fish that never entered the top zone were assigned a maximum latency of 600 seconds (assay duration). BDE-99 doses measured in embryos were 0 (control, CTL), 1.00 (MED), and 2.70 (HI) µg/g dry weight (dw). Sample sizes for males were n = 31 (CTL), 33 (MED), and 32 (HI), and for females, n = 13 (CTL), 17 (MED), and 23 (HI). Within each plot, outcomes (p-values) for statistical tests are shown for the main effects (BDE-99 dose and sex) and their interaction. p<0.05, p<0.01, p<0.001 are indicated by *, **, and ***, respectively. Error bars represent standard error of the mean (SEM).

### Direct Embryonic Exposure: Brain Transcriptome Responses in Adults

Direct embryonic exposure to BDE-99 did not result in persistent changes in brain gene expression in male or female adult fish. In males, WGCNA identified seven co-expressed gene modules, each containing 55 to 249 genes. In females, 12 gene modules ranged from 39 to 537 genes. However, none of these modules showed significant associations with BDE-99 exposure in either sex. Additionally, 12 co-expressed gene modules were identified in the combined (both sexes) analysis, each containing 60-457 genes. Again, none of the modules showed significant associations with BDE-99 exposure. This pattern aligns with the behavioral data showing no effects in males but contrasts with the behavioral findings indicating dose-dependent effects in females.

## Discussion

Our findings demonstrate that relatively brief exposures to BDE-99 during early life lead to lasting behavioral and molecular changes into adulthood, with effects shaped by developmental exposure route, dose, and sex. Developmental exposure to BDE-99, whether via maternal transfer or direct embryonic waterborne exposure, resulted in long-term hyperactivity and reduced anxiety-like behavior, but the specific effects varied. To our knowledge, this is the first study to directly compare these two exposure routes, revealing notable differences in outcomes. The added complexity introduced by exposure route may help explain variability among studies and contributes important nuance to how we predict outcomes based on chemical dose alone. Rather than dose being the sole determinant, exposure-perturbed maternal factors may play an underappreciated role in shaping risk and estimating potential neurodevelopmental damage (Figure 6).

**Figure 6:**
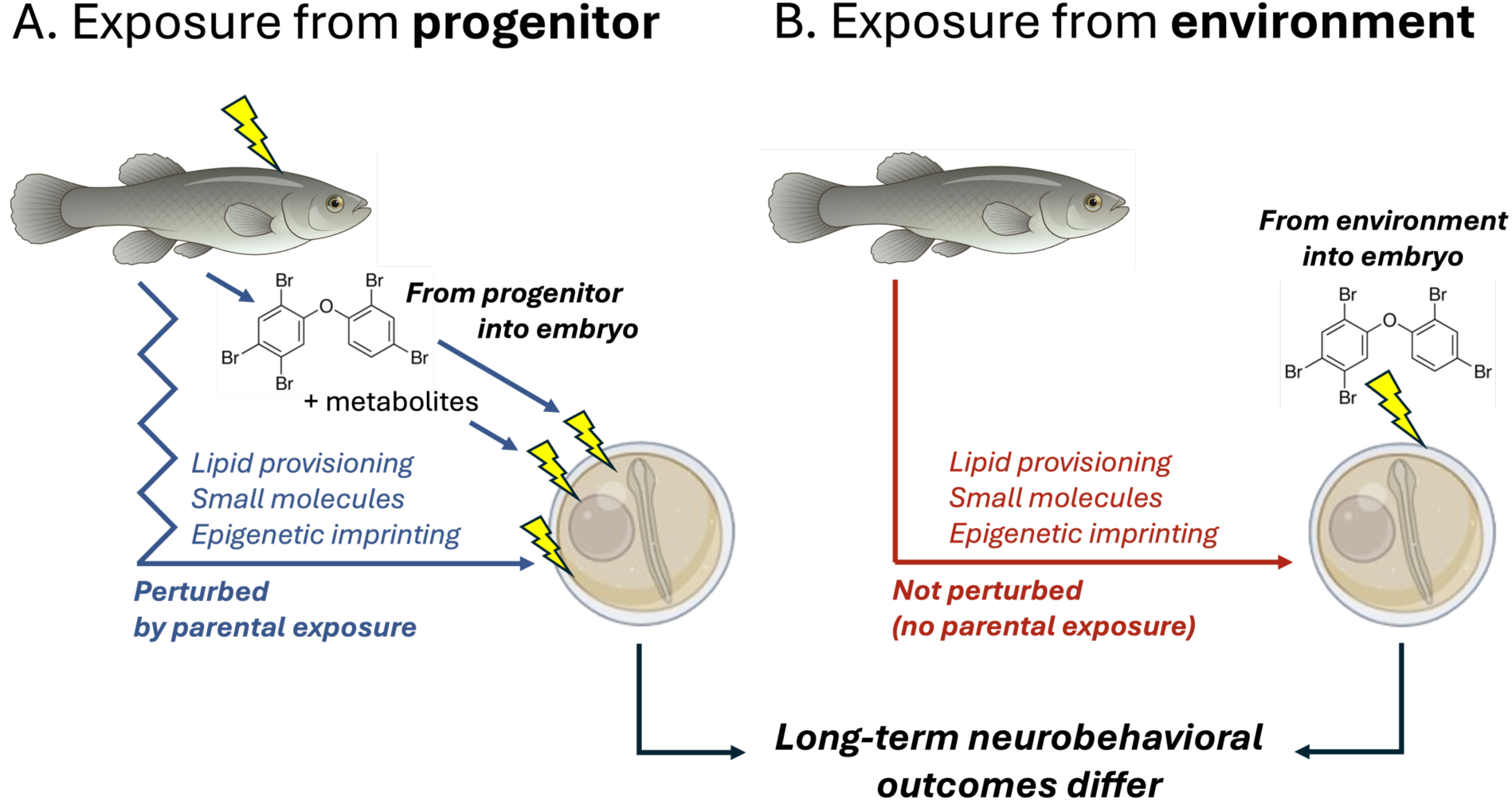
Maternal transfer introduces additional factors that shape offspring responses to PBDE exposure. The route of developmental exposure influences offspring outcomes due to differences in maternal contributions. In progenitor exposure (left panel, blue), maternal transfer of BDE-99 is accompanied by metabolites and additional perturbations, including altered lipid provisioning, small molecules, and epigenetic imprinting. These maternal factors may modify offspring susceptibility to BDE-99 and contribute to long-term behavioral and molecular changes. In contrast, direct environmental exposure (right panel, red) involves only the chemical dose without maternal contributions, resulting in distinct effects. Differences in behavioral and transcriptomic responses between these exposure routes suggest that maternal influences play a critical role in shaping neurodevelopmental outcomes beyond direct chemical exposure alone. While this study examined maternal transfer and environmental exposure separately, both exposure routes occur simultaneously in the wild, potentially exacerbating toxic effects. Fish and embryo illustrations were sourced from BioRender.com.

In the progenitor exposure experiment, behavioral alterations were detected at least in males (Figure 3; females were not assessed due to low sample sizes). In contrast, in the direct embryonic exposure, effects were only evident in females and not in males (Figure 5).

Additionally, progenitor exposure led to long-lasting changes in brain gene expression (Figure 4), while no module-level transcriptomic shifts were detected in males or females from the direct embryonic exposure. Differences in outcomes between exposure routes suggest that maternal factors beyond direct chemical transfer, such as metabolites, hormonal influences, or epigenetic modifications, may contribute to the observed effects (Figure 6). Multigenerational outcomes from BDE-99 exposures also differed between maternal and direct exposure routes in killifish (McNabb-Kelada et al., unpublished). While early-life exposure clearly has lasting consequences, the inconsistencies between behavioral and transcriptomic outcomes and dependencies on sex and maternal effects, plus non-linear dose effects, suggest nuanced and complex dynamics underlying BDE-99-induced neurodevelopmental effects.

BDE-99 exposures during early (embryonic) development led to persistent hyperactivity and reduced anxiety-like behavior in adulthood, which is consistent with the neurobehavioral effects of PBDEs observed in other fish and rodents. Early-life BDE-99 exposure in zebrafish reduced anxiety-related responses in adulthood, even at doses that did not affect larval activity^98^. Early-life exposure to BDE-99 resulted in permanent behavioral disruptions that worsened with age in mice^99^ and caused hyperactivity that persisted into adulthood in rats^100^. Additionally, the non-monotonic dose-response relationship observed, where adult behavioral effects were observed at embryonic doses 0.47, 0.85, and 2.00 µg/g dw, but not at the highest dose (3.50 µg/g), is consistent with multigenerational effects that were non-monotonic in killifish (McNabb-Kelada et al., unpublished). In other mammal and fish species, PBDE exposures also have non-monotonic outcomes for locomotor activity and anxiety-related behaviors^98,101,102^, as well as endocrine and immune disruptions^103–105^. These findings suggest that the relationship between BDE-99 exposure and neurobehavioral effects is non-linear and therefore complex. This pattern should be considered when evaluating PBDE risk, as effects are unlikely to scale linearly with dose.

Persistent behavioral effects of BDE-99 exposure varied by sex and developmental exposure route, contributing complexity to the predictability of exposure-induced behavioral outcomes. In the progenitor exposure experiment, hyperactivity and reduced anxiety-like behavior were evident in adult male offspring, but we could not test for female effects (experimental design limitations). In contrast, in the direct embryonic exposure, only female adults exhibited these behavioral alterations, while males were unaffected. The observed differences between exposure routes suggest that maternal factors beyond chemical transfer, such as hormonal influences and metabolic changes, may play a role in mediating exposure-induced behavioral outcomes (Figure 6). Additionally, maternal transfer of PBDE metabolites, such as hydroxylated PBDEs^106,107^, may have contributed to divergence in behavioral outcomes since directly exposed embryos would not have been exposed to PBDE metabolites during the earliest stages of development. That is, adults (with mature metabolic systems) exposed to PBDEs would have had time to at least partially metabolize the parent compound before transferring parent compound plus metabolites into offspring. In contrast, embryos (with immature metabolism) exposed directly from the water for only a short duration would have had little opportunity to metabolize and thereby be exposed to metabolites, at least during the earliest stages of embryonic development. This difference could help explain our observed divergence in male behavioral outcomes between the progenitor exposure and the direct exposure groups.

At least in the direct exposure experiment, we observed sex-specific persistent behavioral effects, where females were sensitive to early-life BDE-99 exposure but males were insensitive. Evidence for PBDE-induced sex differences in behavior is limited, but some rodent studies suggest that BDE-99 can disrupt neuroendocrine signaling in a sex-specific manner^108–110^.

Additionally, structurally similar persistent organic pollutants such as PCBs impose sex-specific neurotoxic effects, including altered dendritic growth^111,112^ and social behavior^113^. These findings highlight the need for further research to determine whether PBDE-induced sex differences arise through direct endocrine disruption, sex-biased metabolism, or neuroimmune interactions.

Consistent neurobehavioral outcomes following BDE-99 exposure did not always correspond to consistent changes in brain gene expression, suggesting that behavioral alterations may arise through diverse dose-dependent molecular mechanisms or mechanisms other than changes in brain-wide gene expression. In maternally exposed adult fish, significant gene expression differences were observed at both BDE-99 doses (2.00 and 3.50 µg/g) compared to control, yet behavioral effects were only present at 2.00 µg/g. Furthermore, in the direct embryonic exposure experiment, females exhibited behavioral alterations, but without corresponding transcriptomic changes. These inconsistencies may reflect non-genomic mechanisms of PBDE neurotoxicity or limitations of bulk tissue RNA-seq. Comparing differential expression among bulk tissue samples may mask brain region-or cell-type-specific transcriptional changes that are consequential for behavioral phenotypes, where mechanistic insight could be resolved with brain region-specific or single-cell transcriptomics^114^. PBDE neurotoxicity has been shown to involve synaptic and neurotransmitter disruptions that do not necessarily result in widespread transcriptional alterations. In mammals, BDE-99 exposure has been linked to disruptions in glutamate^115^ and nicotinic acetylcholine receptor (nAChR) signaling^116,117^, mitochondrial calcium uptake^118^, and ryanodine receptor (RyR) modulation^119,120^, all of which impair synaptic function. Oxidative stress has also been implicated, as maternal transfer of BDE-99 increased oxidative stress in the hippocampus of rats, coinciding with behavioral alterations^121^. It is plausible that in some exposure conditions, neurobehavioral changes arise primarily through neurotransmitter and synaptic alterations rather than widespread transcriptomic shifts detectable via bulk tissue RNA-seq.

One of the gene expression modules associated with BDE-99 exposure in progenitor-exposed fish was enriched for immune response processes. Immune-related genes were also perturbed by progenitor exposure at the juvenile stage in killifish (McNabb-Kelada et al., unpublished). These observations are consistent with progenitor exposure having a persistent effect on immune gene regulation in their offspring that lasts the duration of their offspring’s life. BDE-99 has been linked to immune system dysregulation, including altered macrophage activity and increased disease susceptibility in salmon^104^. Immune activation has been associated with behavioral changes in zebrafish^122^, highlighting potential neuroimmune interactions.

Additionally, functional enrichment analysis indicated disruptions in protein modification and environmental response pathways, suggesting a broader cellular stress response. While these findings do not establish a direct relationship between gene expression changes and behavioral effects, they provide clues about the potential health consequences of early-life BDE-99 exposure. The lack of consistent alignment between behavioral alterations and transcriptomic changes across exposure groups suggests that multiple, possibly independent mechanisms, including immune activation, neurotransmitter disruptions, and oxidative stress, may be contributing to the observed neurobehavioral effects. Future work should explore whether neuroimmune activation represents a mechanism of PBDE-induced behavioral effects, or rather a secondary response to neuronal dysfunction, and whether a legacy of PBDE exposure influences systemic immunity.

Overall, our findings underscore the complexity of BDE-99-induced developmental neurotoxicity, revealing persistent behavioral effects shaped by exposure route, sex, and dose. Future studies may consider exposure effects on a broader suite of neurobehavioral alterations that could be of consequence for fitness, including cognition, memory, learning, social behavior, and fear/startle responses. There is much opportunity to further define the factors that distinguish outcomes from maternal exposure compared to those from direct exposure, such as exposure effects on maternally transmitted metabolites, small molecules (e.g., lipids, proteins, RNA), and epigenetic imprinting. Additionally, more research is needed to disentangle the roles of neuroimmune interactions, endocrine disruption, and non-genomic mechanisms in mediating PBDE-induced behavioral changes. Illuminating these nuances will be essential for developing more accurate predictive models of PBDE toxicity and more thoroughly estimating risks for ocean health.

## Data Availability

Raw RNA-seq reads are at NCBI (BioProject PRJNA1241279). A matrix of raw gene-level read counts per sample (‘bdeabrn_counts_matrix.tsv’) and RNA-seq analysis scripts are on GitHub (https://github.com/WhiteheadLab/Multigen_BDE_heteroclitus).

## Acknowledgments

Dr. Nadja Brun assisted with setting up the novel tank diving test. Jennifer Roach and Dr. Tony Gill assisted with troubleshooting the BrAD-seq library preparation method. Dr. Joanna Griffiths developed the Snakemake pipeline used to process RNA-seq data. Peyton Delaney assisted with behavioral data analysis. This research was supported by the National Institute of Environmental Health Sciences (T32 ES007059), Fumio Matsumura Memorial Endowment, Jastro-Shields Research Awards, Emmy Werner and Stanley Jacobsen Fellowship, Lewin Family Fellowship, and Schwall Dissertation Fellowship awarded to N.A.M.K.

## Supplemental Methods

We conducted two complementary experiments to test for the long-term effects of early-life exposure to BDE-99 on adult behavior, where we contrasted effects that emerged from two different routes of exposure (Figure 1). In Experiment 1, we investigated the long-term effects of early-life exposure to BDE-99 via maternal transfer by feeding adult progenitors contaminated diets and assessing behavioral, reproductive, and molecular outcomes in their offspring (F1 generation) during adulthood. In Experiment 2, we examined the long-term effects of direct embryonic (waterborne) exposure to BDE-99 by evaluating the behavioral, reproductive, and molecular responses in adulthood. Though many of the methods were identical between experiments, we describe them sequentially below.

### Experiment 1: Progenitor Exposure Chronological Overview

During the breeding season, adult wild-caught killifish were fed either control or BDE-99-contaminated diets for 64 days. This exposure resulted in parental transfer of BDE-99 to their offspring (F1 generation), which were reared under uncontaminated conditions until adulthood (2 years post-hatch) (Figure 1A; the direct embryonic exposure experiment in Figure 1B is described later). At 2 years post-hatch, F1 fish were spawned, and fertilization rates were assessed to evaluate potential reproductive effects of early-life exposure. At 2.5 years post-hatch, novel tank diving tests were conducted to assess anxiety-like behavior, providing insight into long-term behavioral changes. Following behavioral testing, whole brain tissues were flash-frozen and preserved for RNA sequencing (RNA-seq) to examine persistent alterations in gene expression associated with early-life exposure to BDE-99 via maternal transfer.

### Adult Fish Collection and Husbandry

Adult Atlantic killifish (∼10 g wet weight (ww)) were collected using baited traps from Jerusalem, Narragansett, RI, USA (41°22’58.4” N, 71°31’09.6” W) on June 6^th^, 2019, as previously described^1^. This site has historically served as a source of uncontaminated reference killifish, with sediment PCB concentrations ranging from 0 to 4 ng/g dry weight (dw)^2–5^.

Following collection, fish were transported to the US Environmental Protection Agency (EPA) laboratory in Narragansett, where they were housed in large (300 L, 80 gal) tanks with a continuous flow of 23°C, 5 µm-filtered, uncontaminated seawater. They were maintained under a 14:10 h light:dark cycle and fed TetraMin Tropical Flakes (Tetra, Blacksburg, VA, USA) ad libitum. After one month of acclimation, fish were tagged with passive integrated transponder (PIT) tags (Biomark, Boise, ID, USA) for individual identification and distributed into 38-L (10-gal) tanks with a continuous flow of 23°C seawater. Each treatment included three replicate tanks, each containing 14 fish (8 females, 6 males) to encourage breeding. Fish were held under these conditions and fed an uncontaminated diet (described below) for eight days before the experiment began. All procedures using live vertebrate animals at the EPA were conducted following Animal Care and Use Protocols approved by the EPA Institutional Animal Care and Use Committee (IACUC, ACUP # Eco23-03-002, Eco23-11-001, and Eco23-07-001).

### Diet Preparation for Progenitor BDE-99 Exposure

Neat, certified BDE-99 (2,2’,4,4’,5-pentabromodiphenyl ether; >99% pure) was purchased from AccuStandard (Lot no. 27669; New Haven, CT, USA). To prepare a BDE-99 stock solution (2.5 mg/mL), 10 mg of neat BDE-99 was dissolved in 4 mL of acetone (Honeywell Burdick & Jackson, Muskegon, Michigan, USA). For diet preparation, two dosing solutions were made (1.4 mL each): BDE-99_MED and BDE-99_HI. The BDE-99_MED dosing solution for one batch of diet contained 52 µL of the stock solution and 1348 µL of acetone. The BDE-99_HI dosing solution contained 209 µL of the stock solution and 1191 µL acetone. Each batch of diet consisted of 40 g of ground TetraMin Tropical Flakes, amended with either 1.4 mL of acetone (carrier control) or 1.4 mL of the respective BDE-99 dosing solution. The acetone or BDE-99 solution was first added to a 1 L glass bottle, which was rolled to evenly coat the interior and allowed to evaporate completely before adding the ground flakes. The bottle was then sealed and placed on a bottle roller for 7 days to ensure uniform distribution of the chemical treatment.

To supplement the diet, a nutritional mixture was prepared by blending 454 g of frozen chopped spinach, 454 g of Hikari Bio-Pure frozen brine shrimp, 454 g of San Francisco Bay Brand Sally’s frozen krill, 425 g of canned chub mackerel, and 106 g of sardines in oil. The resulting mixture was divided into four equal portions of 450 g each. Each batch of diet was supplemented with 450 g of the nutritional mixture, which was bound together using 20 g of gelatin dissolved in 400 mL hot DI water. The final mixture was thoroughly blended, evenly distributed into aluminum tins on labeled baking sheets, covered with plastic wrap, and left to set overnight. The entire batch was stored frozen and provided enough food for two weeks of feeding across three replicate tanks per treatment, with additional reserves equivalent to one extra tank to account for waste, loss, or samples designated for chemical analysis.

### Progenitor Dietary Exposure to BDE-99

Fish were fed either an acetone-amended control or a BDE-99-contaminated gelatin-based diet at approximately 15% of body weight per day for 64 days. During this period, fish received treatment diets five days per week, while a control diet was provided on the remaining two days. Feeding occurred once daily, with flowing seawater temporarily turned off to ensure consumption. Daily rations were randomly assigned to feeding days, stored at-20°C, and thawed on the day of feeding. The gelatin-based diets were composed of commercial flaked fish food supplemented with small volumes of carrier (acetone) or BDE-99 stock solutions (as described above). Fish were exposed to one of three dietary treatments: control, BDE-99_MED (37.5 ng/g fish ww/day), or BDE-99_HI (150 ng/g fish ww/day). Target internal doses of BDE-99 were 1 and 4 μg/g fish dw, selected based on a preliminary dietary exposure study with *F. heteroclitus* that evaluated a range of BDE-99 concentrations. Dietary exposure was chosen as it better reflects natural exposure routes in wild fish and minimizes handling stress.

### Spawning

The F0 progenitor dietary exposure period (July 9 – September 11, 2019) and spawning days were selected to align with the species’ peak reproductive period, which follows a summer semi-lunar spawning cycle^6^. Fish were manually strip-spawned five times throughout the exposure period to produce F1 offspring that were developmentally exposed to BDE-99 (or control) via maternal transfer. Eggs and milt collected from fish within the same tank were pooled for fertilization but kept separate between tanks. After several hours, embryos were rinsed with clean seawater and incubated at 23°C under a 14:10 h light:dark cycle. At 4 days post-fertilization (dpf), unfertilized eggs were flash-frozen and stored at-80°C for chemical analysis.

In the summer of 2021, 2-year-old F1 fish were spawned across three separate time points to assess persistent reproductive effects of progenitor exposure. Due to low female numbers during the 2021 spawning season, in both F1 control and BDE-99 treatments, males were crossed with external control females collected from the Jerusalem reference site in 2019. Eggs from 20-36 females were pooled, divided into approximately equal groups, and fertilized with pooled milt from males within each treatment. The fertilization rate was recorded at 4 dpf for each spawning time point. A one-way Analysis of Variance (ANOVA) was conducted to analyze fertilization rate (percentage of fertilized eggs per treatment), following confirmation of normality using the Shapiro-Wilk test. Statistical analyses were conducted using GraphPad Prism version 10.4.1 for macOS.

### Chemical Analysis

Analytical-grade hexane and acetone were obtained from Honeywell Burdick & Jackson, while BDE-99 and ^13^C12-BDE-99 (internal standard, IS) were purchased from Cambridge Isotope Laboratories (Tewksbury, MA, USA). Accelerated Solvent Extraction (ASE 200; Dionex Corporation, Sunnyvale, CA, USA) was used to extract BDE-99 from F1 fish egg samples and treatment diets from the progenitor exposure experiment, as well as F0 embryo samples from the direct embryonic exposure experiment^7^. Fish eggs (40-150 mg ww) were ground to a fine powder with sodium sulfate using a mortar and pestle and then transferred to a 2-dram scintillation vial. The samples were spiked with IS, combined with acetone, vortexed for 30 seconds, and centrifuged at 2000 rpm for 5 minutes. Extracts were back-extracted with hexane and concentrated using ultra-high-purity nitrogen in an automatic evaporation system. A 10% aliquot of the extract was reserved for lipid determination, while the remaining portion was treated with concentrated sulfuric acid until the hexane layer was clear.

BDE-99 concentrations were measured using Gas Chromatography-Mass Spectrometry (GC-MS) on an Agilent Technologies (Santa Clara, CA, USA) 6890N GC equipped with a 5973 mass selective detector and an Agilent J&W DB-5ms Ultra Inert analytical column (15 m x 0.25 mm x 0.25 µm). Helium was used as the carrier gas at a constant flow rate of 1.6 mL/min.

Samples were injected in splitless mode with an inlet temperature of 280°C. Ion source and quadrupole temperatures were set at 150°C and 250°C, respectively. Calibration standards for BDE-99 (50-2500 ng/µL) were prepared in heptane, and a seven-point calibration curve was generated for quantification. Data processing was performed using ChemStation software, and BDE-99 concentrations were reported on a wet weight (ww), dry weight (dw), and lipid weight basis. To ensure quality assurance and quality control (QA/QC), procedural blanks, standard reference material (SRM 1947), analytical duplicates, and matrix spikes were routinely processed. Procedural blanks showed no contamination, and the laboratory-measured BDE-99 value was within 10% of the NIST-certified reference value. Analytical duplicates exhibited a relative standard deviation (RSD) of <7%, while matrix spike recoveries ranged from 90% to 104%, confirming method accuracy and precision.

### Rearing and Relocation of Progenitor Exposure F1 Fish

F1 embryos from F0 progenitor exposed and control lineages were incubated in clean seawater in Pyrex glass bowls at 23°C under a 14:10 h light:dark cycle. The seawater was refreshed once a day, and dead embryos were removed. At 7 dpf, embryos were transferred to individual wells in 12-well tissue culture plates (Thermo Fisher Scientific, Waltham, MA, USA) lined with seawater-dampened Restek Cellulose filters (20 mm, Restek, Bellefonte, PA, USA) and maintained at 23°C. Plates were monitored daily to remove any dead embryos. At 10 dpf (late organogenesis, stage 34 ^8^), embryos were screened microscopically for the presence of developmental abnormalities, including pericardial edema, heart abnormalities, cranial or caudal hemorrhaging, and reduced body or head size^9^. There were no treatment effects on developmental abnormalities. At 14 dpf, plates were rocked gently for 1 hr to stimulate hatching. Following this, 3 mL of clean seawater was added to each well containing hatched larvae, which were maintained in their original wells at 23°C. Seawater was renewed every other day, and larvae were fed *Artemia* ad libitum. Larvae were transferred to 9.5 L (2.5 gal) tanks after 14 dph and maintained at 23°C. As fish grew into juveniles, they were gradually transitioned to 19 L (5 gal) and later 38 L (10 gal) tanks to accommodate increasing size and density. Juvenile fish densities were maintained at 3-10 fish per gallon.

In response to the COVID-19 pandemic, all F1 fish (across all treatments) had to be temporarily relocated from the EPA facility in Narragansett, RI to the University of California, Davis for 12 months. Fish from Experiment 2 were subject to the same relocation. Fish were 6-8 months post-hatch at the time of transfer. During this period, they were housed in recirculating artificial seawater systems consisting of 114 L (30 gal) tanks, maintained at densities of 0.4-0.8 fish per gallon. Tanks were aerated and filtered, and room temperature was controlled between 20-21°C under a 12:12 h light:dark cycle. Artificial seawater (20-22 ppt salinity) was prepared using reverse osmosis water and sea salt (Instant Ocean, Crystal Sea, Baltimore, MA, USA).

Water quality was monitored every other day, and 20% water changes were performed weekly to maintain stable conditions. Fish were fed TetraMin Tropical Flakes ad libitum, observed daily, and any mortalities were recorded. The University of California, Davis IACUC approved the protocol for fish maintenance (#22082).

Fish were returned to the EPA facility in April 2021, where they were housed for three months before any experimental endpoints were collected. Out of over 900 fish, only two mortalities occurred during transport to UC Davis, and one during the return to the EPA facility. Upon return, F1 adult fish were housed in flow-through aquaria at 23°C with 5 µm-filtered seawater, maintained at a density of 0.6-1.3 fish per gallon.

### Novel Tank Diving Test

At 2.5 years post-hatch, a novel tank diving test was conducted to assess anxiety-like behavior in adult male fish, as low female numbers across all treatments precluded their inclusion. All testing occurred between 12:00-6:00 pm, with treatment groups evenly distributed across testing times to prevent time-of-day biases. Freshly retrieved flow-through seawater was used for all test tanks and replaced every six trials. The experimental setup followed the protocol reported by Levin et al. 2007, with modifications^10^. Two adjacent rectangular tanks (33.6 cm W x 33.9 cm H x 8.2 cm D) were used, each filled with 16.5 cm of seawater (∼4.5 L). Tank water temperatures were maintained between 21.5 and 23°C. An LED light pad (Huion, Shenzhen, China) was mounted on crossbars above the tanks to provide uniform overhead illumination.

Light levels (180 ± 10 lx) were verified with an LT300 light meter (EXTECH Instruments, Nashua, NH, USA). A Basler acA1300-60gmNIR camera (Basler AG, Ahrensburg, Germany) was positioned 120 cm from the front of the tanks to record the behavioral trials. Video data were captured at 30 frames per second and analyzed without track smoothing using EthoVision XT software (version 11.5.1026, Noldus, Wageningen, Netherlands) for fish tracking and behavioral assessment.

At the start of each trial, two fish were individually placed in 400 mL glass beakers containing 150 mL of seawater and transported to the testing room. Fish were simultaneously released into the novel tank environment and recorded for 10 minutes. Anxiety-like behavior was evaluated using the following metrics: *total distance moved* (cm per 10 min), *duration spent in the top zone* of the tank (percent time), *number of transitions to the top zone*, and *latency to enter the top zone* (seconds). Following behavioral testing, subsets of fish were euthanized using TRICAINE-S (tricaine methanesulfonate, MS-222), and whole brain tissues were flash-frozen in liquid nitrogen and stored at-80°C for molecular (RNA-seq) analysis. Behavioral data were not normally distributed (Shapiro-Wilk test) such that the Kruskal-Wallis test was used to test for treatment effects, followed by Dunn’s multiple comparisons tests to compare the mean rank of each exposure group to the controls. Statistical analyses were conducted with GraphPad Prism version 10.4.1.

### RNA-Seq and Read Count Quantification

Whole brain tissues from adult fish were stored at-80°C until processing. Tissue homogenization was performed using 2.8 mm ceramic beads (Omni International, Kennesaw, GA, USA) on an Omni Bead Ruptor 24 bead mill homogenizer. The resulting lysates were suspended in 750-1300 µL of Lysis Binding Buffer^11^ (volume adjusted based on weight), incubated at room temperature for 10 min, and then centrifuged at 13,000 rpm for 10 min, with the supernatant retained. Messenger RNA (mRNA) was extracted from lysates using oligo (dT)25 beads (Dynabeads^TM^ mRNA DIRECT^TM^ Purification Kit; Invitrogen, Waltham, MA, USA) to enrich for polyadenylated transcripts. The Breath Adapter Directional sequencing (BrAD-seq) method^11^ was conducted to prepare strand-specific RNA-seq libraries, incorporating 13 cycles of PCR enrichment. The fragment priming was accomplished using a random hexamer (GTGACTGGAGTTCAGACGTGTGCTCTTCCGATCTNNNNNNNN, where N represents a random nucleotide). The Qubit^TM^ dsDNA High Sensitivity Quantification Assay Kit (Invitrogen) was used to quantify final libraries, which were uniquely indexed, pooled, and then sequenced across two lanes of NovaSeq X Plus (Illumina, San Diego, CA, USA) as paired-end 150-bp reads by IDseq Inc. (Davis, CA, USA).

Each sample yielded approximately 13 million raw reads. There were three treatment groups represented (control, and 2.00 and 3.50 µg/g BDE-99 progenitor exposure lineages), with 8 biological replicates per treatment, totaling 24 samples. Raw sequencing reads were processed using a Snakemake pipeline developed by Dr. Joanna Griffiths (unpublished; publicly available at https://github.com/JoannaGriffiths/RNASeq-snakemake-pipeline). The pipeline included quality-checking using FastQC^12^ and trimming with fastp^13^ to remove adapter sequences and low-quality bases. Salmon^14^ was used to map processed reads to the *F. heteroclitus* reference genome (GenBank MU-UCD_Fhet_4.1)^15^ for transcript-level quantification. Raw read counts were then summed to the gene level using gene annotation data for gene-level analyses. RNA-seq data will be deposited to the NCBI Sequence Read Archive. All analysis scripts and a matrix of raw gene-level read counts per sample (‘bdeabrn_counts_matrix.tsv’) are available on the Whitehead Lab GitHub site (https://github.com/WhiteheadLab/Multigen_BDE_heteroclitus).

### WGCNA and Functional Enrichment

To identify co-expressed gene modules associated with experimental endpoints, we applied weighted gene correlation network analysis (WGCNA)^16^. This method enables the detection of co-regulated gene clusters across samples by constructing gene networks, or modules, based on expression similarity. First, a variance-stabilizing transformation (VST) in DESeq2 ^17^ was performed to normalize raw gene counts to account for differences in library size. Genes were then filtered based on variance, following WGCNA recommendations, with only the top 10% most variable genes (n=3,164) retained to reduce noise and focus on biologically relevant signals. After preprocessing, pairwise gene expression correlations were computed to construct a signed adjacency matrix. To ensure network robustness, an optimal soft-thresholding power was selected using the scale-free topology criterion. Gene modules, or clusters of co-expressed genes, were then identified using hierarchical clustering combined with dynamic tree cutting. Modules were assigned unique color labels, and eigengene values (the first principal component of module expression) were extracted to summarize expression patterns across samples. Eigengene values were tested for significant associations with treatment (developmental BDE-99 exposure) using one-way ANOVA. To account for multiple comparisons across modules, p-values were adjusted using the False Discovery Rate (FDR) correction (Benjamini-Hochberg method). Modules showing significant associations (adjusted p<0.05) were further analyzed for functional enrichment of Gene Ontology (GO) terms. A Mann-Whitney *U* test^18^ was used to determine whether specific biological processes were overrepresented within gene clusters. All analyses were conducted in R (version 4.4.2)^19^ and RStudio (version 2024.09.1+394).

### Experiment 2: Direct Embryonic Exposure Chronological Overview

Fish embryos from unexposed parents were exposed to either control or BDE-99 via waterborne exposure from 1-7 dpf (Figure 1B). Following exposure, these directly exposed F0 embryos were reared to adulthood under uncontaminated conditions. The same experimental endpoints assessed in the progenitor exposure (Experiment 1) were evaluated for the direct embryonic exposure fish: fertilization success at 2 years post-hatch and adult anxiety-like behavior (novel tank diving test) at 2.5 years post-hatch, including temporary colony relocation during the peak COVID-19 pandemic. After behavioral testing, whole brain tissues were collected and archived for RNA-seq to investigate potential persistent molecular effects of early-life exposure.

### Embryo Collection and Fertilization

Embryos were obtained through mass manual spawning of lab-bred fish, originally derived (one generation previously) from a clean reference population at Scorton Creek, Sandwich, MA (41°45’53.6” N, 70°28’48.0” W)^4,20–22^. Spawning was conducted with four separate breeding tanks, with embryos from each tank kept separate. In each tank, eggs were collected from 20-30 females and pooled, then fertilized with pooled milt from 20-30 males from the same tank. Embryos were incubated at 23°C in Pyrex glass bowls and rinsed with clean seawater several hours after fertilization. Fertilization success was checked at 1 dpf to ensure that only fertilized embryos proceeded to the dosing.

### Direct Embryonic Exposure to BDE-99

A BDE-99 stock solution (2.5 mg/mL) was made by mixing 10 mg of neat BDE-99 (Accustandard) into 4 mL of acetone (Honeywell Burdick & Jackson). For each exposure experiment, fertilized embryos from the four breeding groups were equally distributed among three treatment conditions (control and two BDE-99 concentrations) and further subdivided into two replicates per treatment. This design resulted in a total of 24 exposure jars (4 breeding groups x 3 treatments x 2 replicates per treatment). Embryos were placed in glass jars at a density of 1 embryo per 2 mL of seawater and mass exposed from 1 to 7 dpf. Exposure solutions contained a 1% dosing solution, consisting of either carrier (acetone, control) or BDE-99 in acetone at final concentrations of 6.2 ng/mL (MED) and 26 ng/mL (HI). Jars were incubated at 23°C, swirled daily, and checked for mortality, with dead embryos removed as needed. At 7 dpf, embryos were transferred to individual wells in 12-well plates lined with seawater-dampened Restek Cellulose filters (20 mm) and reared following the protocol detailed in the *Embryo-Larval Rearing and Early-Life Endpoints* section. The entire exposure experiment was repeated across three separate spawning events, aligned with the species’ semi-lunar spawning cycle. Due to low sample sizes in one of the breeding groups, embryos from that group in the third exposure experiment were archived for chemical analysis (detailed in the *Chemical Analysis* section) to determine representative doses and BDE-99 uptake in the embryos.

### Rearing and Relocation of Direct Embryonic Exposure F0 Fish

Fish from the direct embryonic exposure experiment were reared under the same conditions as previously described in the *Rearing and Relocation of Progenitor Exposure F1 Fish* section. Juvenile density was maintained at 5.5-12.5 fish per gallon. At 6-8 months post-hatch, all direct exposure F0 fish (all treatments) were temporarily relocated to UC Davis. All rearing conditions were the same as previously described, except that tank density was 0.5-1 fish per gallon. Upon their return to the EPA facility, fish were housed at a density of 0.7-1.3 fish per gallon for 3 months before experimental endpoints were assessed.

### Experimental Endpoints

To assess the long-term reproductive effects of developmental BDE-99 exposure, males and females were crossed within their prior control or BDE-99 treatments. Eggs were collected via manual strip-spawning, pooled within each replicate group, and fertilized with pooled milt from males in a different replicate group within the same exposure condition. Fertilization success was evaluated at 4 dpf. A one-way ANOVA was conducted to analyze fertilization rate (percentage of fertilized eggs per treatment) following confirmation of normality using the Shapiro-Wilk test.

At 2.5 years post-hatch, the novel tank diving test was conducted to assess anxiety-like behavior in both male and female fish, following the same methodology that was described for Experiment 1 (in the *Novel Tank Diving Test* section above). A generalized linear model (GLM) with least squares regression was used to test the effects of sex and BDE-99 dose on behavioral outcomes. The model included the main effects of sex (categorical: male, female) and dose (categorical: CTL, MED, and HI (0, 1.0, and 2.7 µg/g)), as well as their interaction. GLMs were chosen because data did not meet normality and homoscedasticity assumptions, assessed using Shapiro-Wilk normality tests and Spearman’s test for heteroscedasticity. To correct for heteroscedasticity, inverse weighting (1/Y) was applied. All statistical analyses were performed in GraphPad Prism version 10.4.1.

### RNA-Seq and WGCNA

Whole brain tissues from the direct embryonic exposure adult fish were processed as described in the *RNA-Seq and Read Count Quantification* section. Strand-specific RNA-seq libraries were prepared and sequenced together with the progenitor exposure libraries, following the same methodology and specifications. Each sample yielded approximately 12 million raw reads. The dataset included two treatments (CTL and HI), with both combined and separate analyses for males and females. For males, there were 2-6 replicates each from 3 breeding groups per treatment, totaling 27 samples. For females, there were 3-6 replicates each from 3 breeding groups per treatment, totaling 30 samples. Raw sequencing reads were quality-checked and processed as previously described. WGCNA was performed separately for males and females, using the same parameters and workflow outlined in the *WGCNA and Functional Enrichment* section.

